# Forest fragmentation and impacts of intensive agriculture: responses from functional groups of the tree community

**DOI:** 10.1101/546796

**Authors:** Juliana C. Tenius Ribeiro, André Felippe Nunes-Freitas, Mariella Camardelli Uzêda

**Affiliations:** Department of Environmental Sciences, Forest Institute, Federal Rural University of Rio de Janeiro, Seropedica, Rio de Janeiro, Brazil; National Center for Research in Agrobiology, Brazilian Agricultural Research Corporation, Seropedica, Rio de Janeiro, Brazil

**Author notes:** Laboratory of Forest Ecology and Plant Biology, Department of Environmental Sciences, Forest Institute, Federal Rural University of Rio de Janeiro, Seropedica, Rio de Janeiro, Brazil (André Felippe Nunes-Freitas). National Center for Research in Agrobiology, Brazilian Agricultural Research Corporation, Seropedica, Rio de Janeiro, Brazil (Mariella Camardelli Uzêda).

## Abstract

Agricultural landscapes are seen as areas of extreme importance for studying and developing strategies that integrate biodiversity conservation and ecosystem services with food production. The main strategies for intensifying agriculture are based on conventional practices of frequently using agricultural inputs for fertilization and correction of soil pH. Some studies show that these practices generate impacts on nearby forest fragments through soil contamination, causing an increase in nutrient content. The objective of this study was to identify the impacts on the functional groups of sciophilous and heliophilous species of a tree community of 14 forest fragments near agricultural areas under conventional practices, and raised the hypothesis that the higher the fertility of forest fragments adjacent to intensive agriculture modifies the floristic composition of the tree community. The floristic composition of fragments close to agricultural areas are more similar to each other and the General Linear Model (GLM) results show a clear influence of the intensive farming environment on the richness and abundance of the two functional groups in the forest fragments, directly benefiting the abundance of heliophilous species which are also benefited by the greater declivity and smaller fragment area, while the abundance of sciophytes is negatively correlated with these last two variables. The increase of calcium content is beneficial for the richness of heliophilous species, while the increase in phosphorus content influences a reduction in the richness of sciophyte species, which also respond strongly to the isolation between fragments. The results indicate a dominance trend of pioneer species in nutritionally enriched soils, evidencing that the intense adoption of inputs in cultivated areas causes concrete impacts on the diversity of the tree community of forest fragments, being more determinant for the species richness than the size of the fragments.

## Introduction

The loss of habitat promoted by the anthropic transformation of land use is currently the cause of greater impacts to biodiversity [1]. In addition to the loss of habitat through reduced natural vegetation cover, the creation of small isolated fragments in an altered landscape brings severe consequences to the ecological interactions necessary to maintain biodiversity and ecosystem functions [2-4]. With the global demand for increased food production, agriculture is in increasing expansion [5,6], mainly in tropical countries [7], and is one of the main activities causing deforestation worldwide [8], occupying almost 40% of the soil throughout the planet [9].

For this reason, agricultural landscapes are seen today as areas of extreme importance for studying and developing strategies that integrate biodiversity conservation and ecosystem services with food production [6,10]. Thus, the discussion on the conservation of forest fragments in landscapes with a predominance of agricultural activity raises the need to develop production models that not only take into account greater production efficiency, but also the externalities that the different productive systems imply on the biodiversity of a landscape [6, 10-14].

The main strategies for agriculture intensification are based on technological packages that include conventional practices and the frequent use of agricultural inputs for fertilization and correcting soil pH [5,6]. Some studies show that the intensification of these practices in agricultural fields generates impacts on nearby forest fragments through soil contamination by fertilizers, causing increased nutrient content in these areas [15-18]. These changes in fertility levels of soil fragments can have significant impacts on the floristic composition due to the great correlation between vegetation and soil chemical characteristics [19-21].

Some nutrients are limiting to the growth of trees in forest environments [22-25], and the continuous increase of fertility levels in forest fragment soils may lead to alterations in soil chemical relationships [24,26] and possible species losses [27]. Some studies suggest that some species are more efficient in using nutrient surpluses, with an increase in growth rates [24,25]. Due to this, some understory species may present dominance in forest soils enriched with nutrients such as calcium and nitrogen, as shown in some studies in temperate climates [16,29,30], causing long-term species replacement [30]. Thus, continuous additions of nutrients to the soils of forest ecosystems over a long period may alter the functional diversity and composition of plant species of a community [22].

Thus, the present study aimed to identify the impacts on the functional groups of sciophilous and heliophilous species in the tree community of forest fragments near agriculture areas with conventional practices of intensive fertilizer use. The work hypothesized that the higher fertility of forest fragment soils adjacent to intensive agriculture modifies the floristic composition of the tree community, adopting the premise that there are nutrients added into the soils of these fragments.

Studies which aim to observe vegetation patterns with isolated factors such as the chemical characteristics of the soil have limitations due to the great correlation between the soil parameters and the vegetation itself to several other factors [20]. Therefore, in order to identify the impact of a change in the fertility levels within forest fragments regarding the floristic composition of these sites, the vegetation correlation with local factors related to the soil also need to be evaluated, such as slope, soil size and canopy opening, as well as factors related to forest fragmentation such as size, isolation and shape of the fragments, which directly affect the floristic composition. Thus, this study was guided by the following questions:

i. Do forest fragments with intensive farming environments present differences in floristic composition of species?
ii. Does the soil fertility influence the tree species composition?
iii. Which variables influence the species abundance and richness in the forest fragments with different types of use around their environment?

## Materials and Methods

### Site description

The study was carried out in the Guapi-Macacu Basin in the state of Rio de Janeiro, located east of Guanabara Bay. The predominant climate in the region is humid tropical Af according to the Köppen classification (1948). The average precipitation varies between 1,300 and 2,200 mm and the temperature between 14 and 27 °C, presenting an average of 21.1 °C [30]. The vegetation of the region is inserted in the Atlantic Forest (sensu stricto), with the Dense Ombrophilous Lowland Forest phytophysiognomy being predominant [31]. The forest cover of the basin occupies 42.4% of the territory and is divided into larger and continuous fragments in areas of higher elevations, while hills and hillocks are located in the lowlands with smaller and more dispersed fragments in the landscape [30].

The study was developed in 14 forest fragments dispersed by the Basin, with an environment predominantly consisting of agriculture and livestock. The fragments are surrounded by different soil uses, predominantly pastures and crops (43.6% and 4.8% of the territory, respectively) (Fig 1). Agriculture herein will be treated as intensive crop (IC), with high use of agricultural inputs and soil tillage, and livestock treated as extensive crop (EC). The forest fragments of this study vary in size from 8 to 260 hectares and portions of continuous forests belonging to the Three Peaks State Park (*Parque Estadual dos Três Picos*).

**Fig 1.**
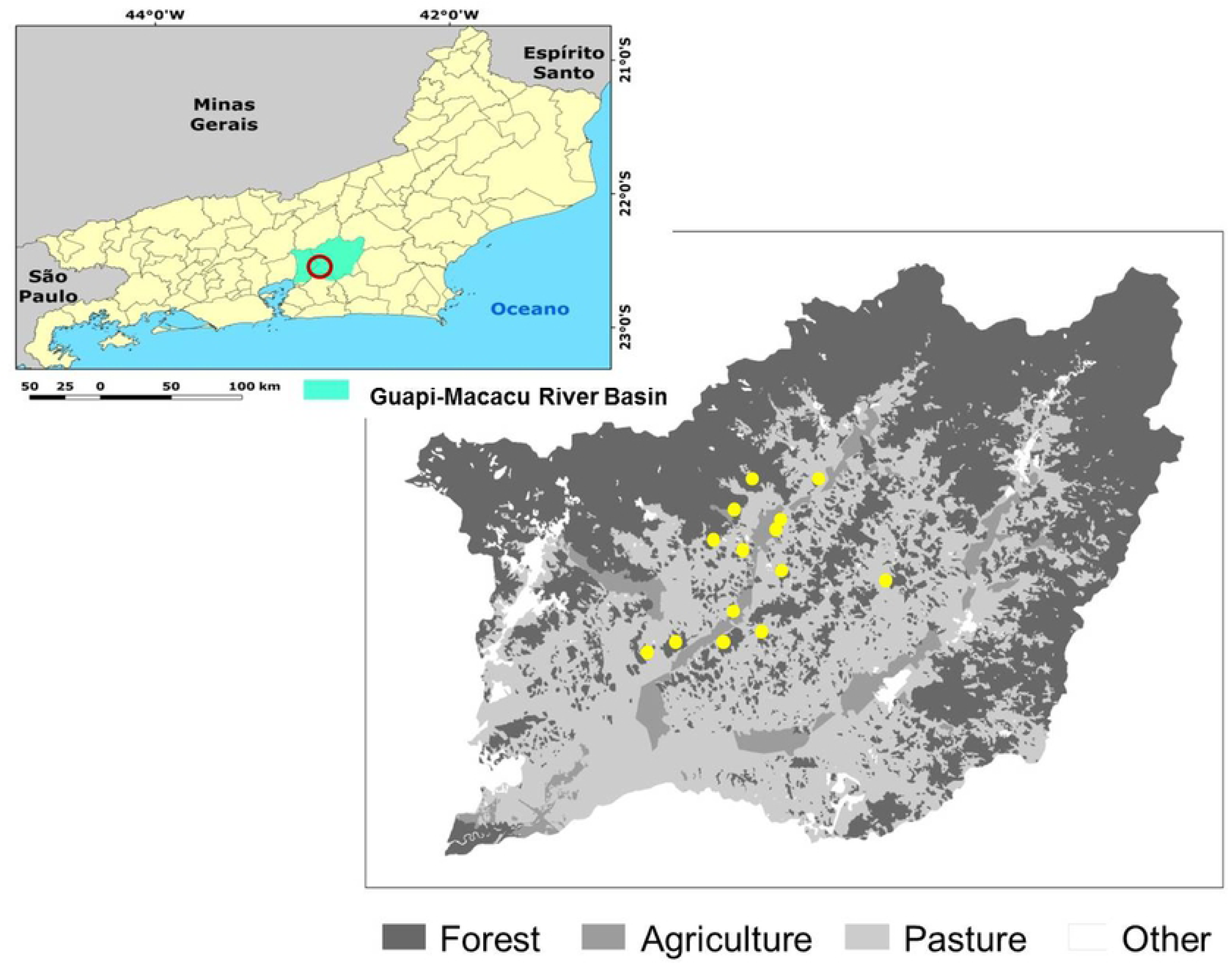
Location of the studied fragments and land use classes in the Guapi-Macacu River Basin, Rio de Janeiro, Brazil (FIDALGO et al., 2008).

Intensive use areas are characterized by annual corn (*Zea mays*) rotations with manioc cultivation (*Manihot esculenta*), with high frequency of soil rotation through plowing followed by harvesting and use of fertilizers and limestone. An average of 2000 kg.ha^−1^ limestone and 60 kg^−1^ of 4-14-8 NPK fertilizer are added for the corn cultivation. Some proprietors use organic compost (manure) applied as cover as a complement to chemical fertilization. Agrochemicals are also frequently used to control pests such as caterpillars (*Spodoptera frugiperda*). Next, a new plowing and harrowing is carried out for planting the manioc roots with an influence of the residual fertilization effect for corn. Management activities are almost non-existent in pasture areas (*Brachiaria brizantha*), with only general grazing occurring. The maximum stocking density of cattle is on average 1 head.ha^−1^.

The criteria described by [32] were adopted as reference to evaluate the effect of the soil use on the native vegetation areas. These authors establish that the anthropic area of the environment must have a minimum size of 100 m in length and the same area of width in contact with the fragment edge. Thus, the selected fragments would have a border in direct contact with the agroecosystems, with 7 fragments in contact with those of extensive use (pasture/livestock) and 7 in contact with intensive use (agricultural cultivation). All those classified as IC have a continuous usage history of adjacent agroecosystems for at least 10 years.

### Sample

The selected forest fragments are characterized by a sloping hillside relief, and thus three different strata were established for sampling, with the sole objective of sampling the differences in slope and distances from the edge of the fragment to the production areas. The fragments were stratified as follows: i) 1^st^ third - region closer to the agroecosystem, which suffers greater anthropic influence; ii) 2^nd^ third – the region next to the 1^st^ third, generally with great slope; iii) Final third - top hill region, with lower slopes. Three plots of 50 x 5 m (250 m^2^) were allocated in each stratum, which totals an area of 2,250 m^2^ per fragment. The plots were systematically delimited in each stratum, facing the fragment directed towards the agroecosystem, with 10 meters of distance between them in the horizontal and 10 meters in the vertical. Three plots were also allocated in the surrounding agricultural areas for soil sampling, following the same methodology for allocating the plots within the fragments.

### Floristic and phytosociological survey

A forest inventory was carried out in March 2016 as assistance to a botanist. All individuals with chest circumference (CBH) greater than or equal to 15 cm were measured and identified in all the plots. When possible, the identification was carried out in the field and, when necessary, botanical material collections were carried out for further identification. The taxonomic determinations were carried out by consulting the Herbariums of the Research Institute of the Botanical Garden of Rio de Janeiro and the Department of Botany (RBR) of UFRRJ (Federal Rural University of Rio de Janeiro), by specialized bibliography, or by specialists. Part of the floristic survey had already been carried out in a previous study [18] using the same methodology used in this study. These data were also collected for conducting this work.

The species were classified into two ecological groups according to Swaine and Whitmore^33^: pioneer or heliophilous species, which require direct light for seed germination, or those generally classified as pioneers and early secondary; and the group of non-pioneer/late or sciophilous species which can germinate and develop under the shade, being found under the canopy and also in open environments [34]. Studies for the state of Rio de Janeiro were used for these classifications in order to avoid variations in classifications [35-38]. Non-classified species (NC),did not fall into any of the categories due to lack of information.

The calculations of the community phytosociological parameters were performed according to Mueller-Dombois & Ellenberg^39^ by calculating the absolute and relative parameters of species density, dominance and frequency in order to obtain the Importance Value (IV) of each species, which consists of the sum of the relative values mentioned above (Table S1).

### Soil fertility and granulometry

Soil samples were collected between February and March 2016 in order to analyze the relation of the tree species with the fertility characteristics of soil of the forest fragments, in which three simple soil samples were collected per plot at a depth of 0-5 cm with the aid of a metal probe. Samples were collected at the ends (0; 50 m) and center (25 m) of each plot, forming a sample composed of soil for each plot. Each composite sample was placed in a plastic bag and identified for transport to the laboratory, where each soil sample was air dried. After drying, they were discharged and sieved using an 8 mm sieve to remove coarse material. The samples were sent for chemical analysis in the Nutrient Cycle Laboratory of EMBRAPA Agrobiology, which follows the methods recommended by EMBRAPA^40^. The chemical characteristics analyzed were pH in water, calcium (cmolc.dm^−3^), magnesium (cmolc.dm^−3^), potassium (mg.L^−1^), phosphorus (mg.L^−1^), carbon (dag.Kg^−1^) and nitrogen (dag.Kg^−1^). The granulometric analysis of the samples was carried out at the Soil Physics Laboratory of the *UFRRJ* Soils Department, using the Pipette Method [40], quantifying the sand, silt and clay fractions expressed as percentage.

### Canopy Opening

Hemispheric photographs were taken for estimating the canopy cover of each fragment, from which it was possible to indirectly calculate the canopy cover and light input in the plots [41]. The photographs were taken using a digital camera attached to a ‘fish eye’ lens positioned at a distance of 1.5 m from the ground using a tripod. The tripod was always positioned to the north with the help of a compass in order to maintain standardization of the photographs. Afterwards, each photo underwent treatment with the purpose to quantify white (the points related to the open sky) and black (referring to the vegetation) pixels, which was performed using the Gap Light Analyzer (GLA), Version 2.0 program [42]. Analysis of the photographs enabled estimating the canopy opening, direct light input and diffuse light. Direct light input was used as a measure of canopy opening for the statistical analysis of the present study due to the high correlation of these parameters.

### Slope

The central point in each plot (25 m) was measured with the aid of a digital clinometer for evaluating the terrain slope.

### Obtaining and evaluating landscape indexes

In order to understand the relationships between the structural variables of the landscape and the tree community composition in the fragments, landscape metrics were used according to the procedures used in Uzêda et al.^18^ obtained with the assistance of the researcher Dr. Elaine Fidalgo from the Geomatics Nucleus (NGEO) of Embrapa Soils. The following measures were used: area (area), which refers to the fragment size; perimeter/area ratio (PARA), which is an indicator of the fragment shape, and therefore related to the amount of existing border; and Euclidean distance of the nearest neighbor (ENN), which is an indicator of fragment isolation. The agricultural bordering limit (limagri) was also obtained, which is an indicator of the border perimeter percentage of the forest fragment which is directly in contact with the agricultural area. All types of land use along the fragment perimeter that made direct contact with its border which were delimited in the map and coverage used were observed for this calculation.

The metrics were calculated using the land use and coverage map of the Guapi-Macacu and Caceribu river basins in 2007, at a scale of 1:50,000 [43]. This map was elaborated based on the image classification of the TM-Landsat 5 sensor, from June to August 2007, with a resolution of 30 meters. The original map was cut and the data of the fragments were specialized in “raster” format with resolution of 30 meters. Thus, the metrics were calculated from this through the Fragstats program [44]. Then, the proportion of their boundaries to be found in line with the different types of land use was calculated. Because the studied fragments are only surrounded by pasture and agriculture, and therefore the percentages of these two limit types total 100, it was decided to only use the limit percentage with agriculture, thus avoiding highly correlated answers [18].

## Data analysis

In order to identify the similarity of the structure and composition of the tree community among the fragments, non-metric multidimensional scaling (NMDS) with Bray-Curtis similarity index [45] was used to verify clustering trends. Therefore, a matrix with the Importance Value (IV) of the species of each fragment (Table S2) was used. NMDS construction was performed in the statistical R program using the “vegan” packages, version 2.3.0, and the “labdsv” version 1.7.0 [46].

Generalized linear models (GLMs) were constructed and tested to identify the variables that best explain the tree species composition pattern in the fragments. Therefore, in order to construct these predictive models, the richness and abundance of the ecological groups (pioneer and late) were considered as dependent variables and the levels of carbon, nitrogen, phosphorus, potassium, calcium and magnesium (C, N, K, Ca and Mg, respectively), clay percentage in the soil (Arg), border limit percentage with agriculture (limagri), Euclidean distance of the nearest neighbor (ENN), fragment size in hectares (area), perimeter-area ratio (PARA), canopy opening (abos) and slope as independent variables (Table S3). We used the data of abundance of each species and average data of the independent variables in each stratum of each fragment as inputs. Next, 19 possible models were tested for the abundance of heliophytes and 28 models for the abundance of sciophytes in total. Then, 33 models were tested for the richness of heliophilous species, and 27 models for the richness of the sciophilous species (Table S4).

Considering the Gaussian data distribution, the models were tested and selected through the second order Akaike criteria (AICc) (Burnham & Anderson, 2002). Models with ΔAICc values smaller than two (Δ(ΔAICc <2) and of higher AICc weight (AICcWi) were selected. The evaluation of adjusting the parameters of the selected models was performed by a chi-square test (χ^2^). The analyzes of the models were carried out in the R statistical software program [46] using the packages “bbmle”, version 1.0.16, and “MuMIn”, version 1.15.1.

## Results

A total of 4922 individuals were sampled from trees and palm trees, and 371 morphospecies were identified in 58 botanical families for the 14 forest fragments (total sampling area of 3.15 ha) studied in the Guapi-Macacu Basin. Of the morphospecies, 242 (66.58% of the total) were identified at the specific level, 61 (16.44%) at the genus level, 36 (9.70%) at the family level and 27 (7.28%) remained undetermined (Table S1). The richness of the fragments ranged from 50 species (ICp1) to 109 species (ECg1).

The non-metric multidimensional scaling (NMDS) of the tree species community composition of the fragments (Fig. 2) reflected the tendency of the sites belonging to fragments adjacent to the EC areas to be grouped, predominantly in the lower quadrants of the graph, whereas the IC fragments are concentrated in the quadrants.

**Figure 2.**
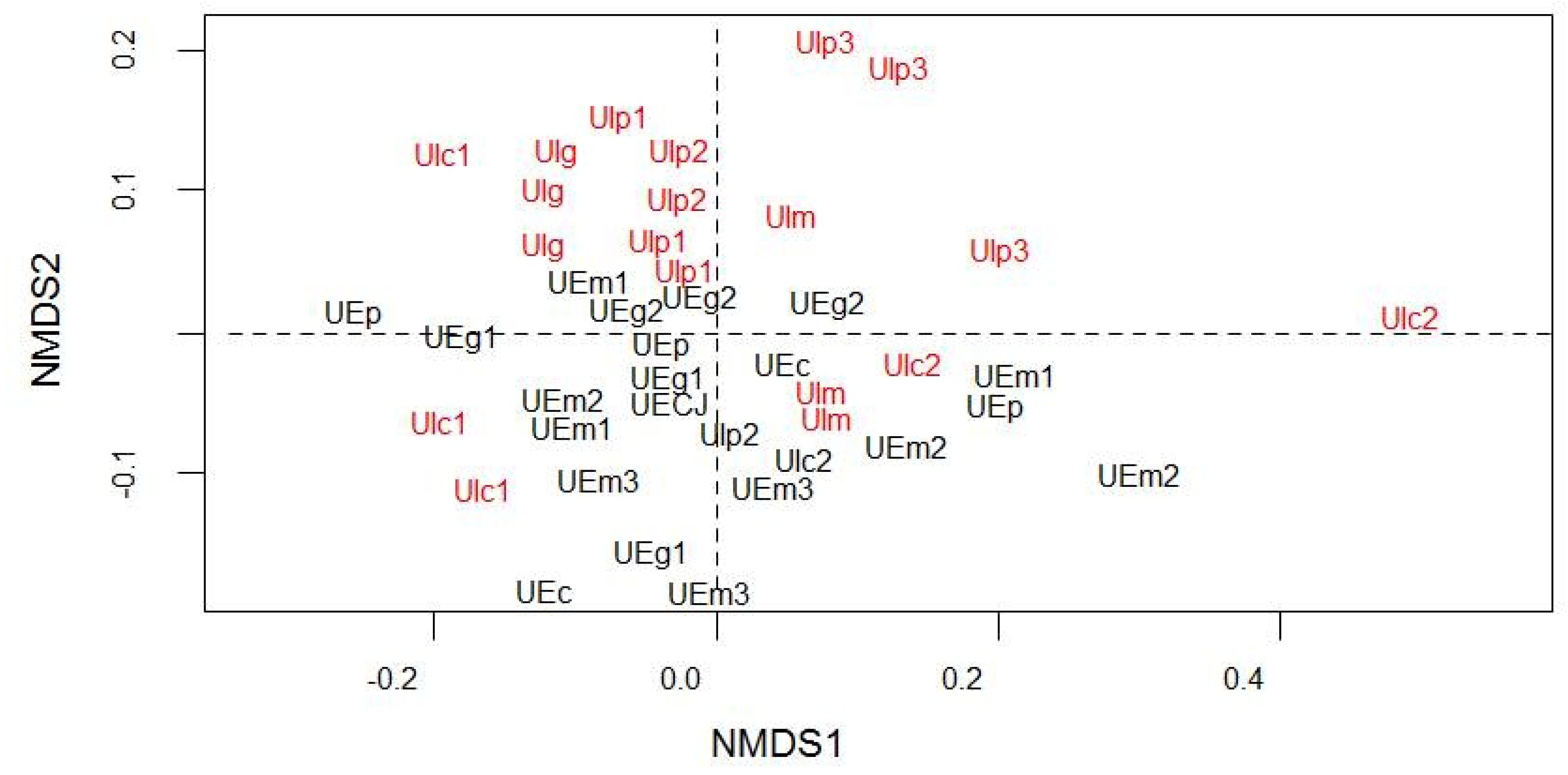
Multidimensional non-metric scaling of the structure and composition of the tree species community in fragments of small (p), medium (m), large (g) and continuous (c) sizes, adjacent to intensive (IC) and extensive (EC) agricultural environments. Fragments neighboring agricultural fields (IC) are in red and fragments neighboring grassland (EC) are in black. *Stress* = 0.11; R^2^= 0.966.

Regarding the abundance of the heliophilous species, GLM indicated that the greater extent of direct limit with agriculture (p < 0.001) and higher slope (p < 0.01) positively affected the number of individuals, while the higher magnesium levels (p < 0.05) and the increase in the size of the fragments (p < 0.05) had a negative relation with the number of individuals in this functional group. Three explanatory models (Table 1) were selected for the abundance of heliophiles, in which the variables of agricultural limit, fragment size, slope and calcium and magnesium contents were included, and the latter two factors were not significant by the chi-square test. The predictive model with the highest weight (AICcWi = 0.3167) showed the agricultural limit (p < 0.001), the slope (p < 0.01) and the fragment size (p < 0.1) as positive explanatory variables. The second model (AICcWi = 0.1686) also presented the magnesium content (p < 0.05) as explanatory, negatively interacting in the number of tree individuals in the heliophilous group.

**Table 1:** Results of the selected GLM models (ΔAICc < 2) to explain the abundance of heliophilous species in the forest fragments of the Guapi-Macacu Basin. The independent variables, the term used in the model, their respective coefficients and their significance by the Chi-square test (χ^2^) are specified below the models.

| Model/Parameters | Term | Coefficient | $\chi^2$ |
| --- | --- | --- | --- |
| Heliophilous = $\beta_0 + \beta_1 \text{ limagri} + \beta_2 \text{ area} + \beta_3 \text{ slope}$ | | $\Delta AICc = 0$ | $AICcWi = 0.3167$ |
| Intercept | Intercept | 145.900 |  |
| Agri limit | limagri | 576.40 | 0.0001708 *** |
| Fragment size | area | -0.41 | 0.0511857 . |
| Slope | slope | 1.478 | 0.0052936 ** |
| Heliophilous = $\beta_0 + \beta_1 \text{ Ca} + \beta_2 \text{ Mg} + \beta_3 \text{ limagri} + \beta_4 \text{ area} + \beta_5 \text{ slope}$ | | $\Delta AICc = 1.1$ | $AICcWi = 0.2113$ |
| Intercept | Intercept | 165.000 |  |
| Calcium content | Ca | 56.410 | 0.110645 |
| Magnesium content | Mg | -138100 | 0.046768 * |
| Agri limit | limagri | 597.90 | 6.792e-05 *** |
| Fragment size | area | -0.56 | 0.010340 * |
| Slope | slope | 1.623 | 0.003206 ** |
| Heliophilous = $\beta_0 + \beta_1 \text{ Mg} + \beta_2 \text{ limagri} + \beta_3 \text{ area} + \beta_4 \text{ slope}$ | | $\Delta\text{AICc} = 1.3$ | $\text{AICcWi} = 0.1686$ |
| Intercept | Intercept | 15.4200 |  |
| Magnesium content | Mg | -60910 | 0.22489 |
| Agri limit | limagri | 605.70 | 8.551e-05 *** |
| Fragment size | area | -0.49 | 0.025151 * |
| Slope | slope | 1721 | 0.002357 ** |

\*\*\* significant at 0.1%, \*\* 1%, \* 5% and . 10% probability.

Next, four explanatory models were selected for the abundance of the sciophilous species (Table 2). Only the increase in the area of the fragments presented a positive relation with the abundance of these late species (p < 0.05), while the slope was the only factor that presented a negative relation with the abundance of individuals in the sciophilous group (p < 0.05). The agricultural limit, phosphorus and calcium content variables were not significant by the chi-square test in these models.

**Table 2:** Results of the selected GLM models (ΔAICc < 2) to explain the abundance of sciophilous species in the forest fragments of the Guapi-Macacu Basin. The independent variables, the term used in the model, their respective coefficients and their significance by the Chi-square test (χ^2^) are specified below the models.

| Model/Parameters | Term | Coefficient | $\chi^2$ |
| --- | --- | --- | --- |
| Sciophilous = $\beta_0 + \beta_1 \text{ P} + \beta_2 \text{ area} + \beta_3 \text{ slope}$ | | $\Delta\text{AICc} = 0$ | $\text{AICcWi} = 0.2146$ |
| Intercept | Intercept | 7.97E+08 |  |
| Phosphorous content | P | -4.46E+07 | 0.12067 |
| Fragment size | area | 3.39E-04 | 0.04164 * |
| Slope | slope | -0.78 | 0.05859 . |
| Sciophilous = $\beta_0 + \beta_1 \text{ limagri} + \beta_2 \text{ area} + \beta_3 \text{ slope}$ | | $\Delta\text{AICc} = 0.6$ | $\text{AICcWi} = 0.1616$ |
| Intercept | Intercept | 6.30E+08 |  |
| Agri limit | limagri | -0.15 | 0.17479 |
| Fragment size | area | 0.00 | 0.02455 * |
| Slope | slope | -0.89 | 0.02994 * |
| Sciophilous = $\beta_0 + \beta_1 \text{ Ca} + \beta_2 \text{ area} + \beta_3 \text{ slope}$ | | $\Delta\text{AICc} = 1.8$ | $\text{AICcWi} = 0.0857$ |
| Intercept | Intercept | 6.08E+04 |  |
| Calcium content | Ca | -1.56E+04 | 0.44894 |
| Fragment size | area | 0.3878 | 0.02050 * |
| Slope | slope | -788.1 | 0.07184 . |
| Sciophilous = $\beta_0 + \beta_1 P + \beta_2 \text{ limagri} + \beta_3 \text{ area} + \beta_4 \text{ slope}$ | | $\Delta\text{AICc} = 2.0$ | $\text{AICcWi} = 0.0792$ |
| Intercept | Intercept | 7.71E+08 |  |
| Phosphorous content | P | -3.48E+07 | 0.25275 |
| Agri limit | limagri | -1.01E-01 | 0.38941 |
| Fragment size | area | 3.34E-04 | 0.04283 * |
| Slope | slope | -7.94E-01 | 0.05170 . |

\*\*\* significant at 0.1%, \*\* 1%, \* 5% and . 10% probability.

Two predictive models were selected for the richness of the heliophilous species (Table 3). Both the higher weight model (AICcWi = 0.3066) and the lower weight model (AICcWi = 0.1768) indicated calcium (p < 0.01) and soil clay levels (p < 0.001) as positive, and the greater opening of the canopy (p < 0.001) as negative.

**Table 3:** Results of the selected GLM models (ΔAICc < 2) to explain the richness of heliophilous species in the forest fragments of the Guapi-Macacu Basin. The independent variables, the term used in the model, their respective coefficients and their significance by the Chi-square test (χ^2^) are specified below the models.

| Parameters | Term | Coefficient | $\chi^2$ |
| --- | --- | --- | --- |
| Heliophilous = $\beta_0 + \beta_1 \text{ Ca} + \beta_2 \text{ Cano} + \beta_3 \text{ Clay}$ | | $\Delta\text{AICc} = 0$ | $\text{AICcWi} = 0.3066$ |
| Intercept | Intercept | 15.52 |  |
| Calcium content | Ca | 9.04 | 0.0011481 ** |
| Canopy opening | Cano | -0.2094 | 0.0002689 *** |
| Clay content | Clay | 0.2049 | 0.0005280 *** |
| Heliophilous = $\beta_0 + \beta_1 \text{ Ca} + \beta_2 \text{ Clay} + \beta_3 \text{ ENN} + \beta_4 \text{ Cano}$ | | $\Delta\text{AICc} = 1.1$ | $\text{AICcWi} = 0.1768$ |
| Intercept | Intercept | 14.95 |  |
| Calcium content | Ca | 8.59 | 0.0016990 ** |
| Clay content | Clay | 0.20323 | 0.0004671 *** |
| Isolation | ENN | 0.01462 | 0.2013926 |
| Canopy opening | Cano | -0.24649 | 0.0001455 *** |
| Heliophilous = $\beta_0 + \beta_1 \text{ Ca} + \beta_2 \text{ ENN} + \beta_3 \text{ Cano} + \beta_4 \text{ Clay}$ | | $\Delta\text{AICc} = 1.1$ | $\text{AICcWi} = 0.1768$ |
| Intercept | Intercept | 14.95 |  |
| Calcium content | Ca | 8.59 | 0.0016990 ** |
| Isolation | ENN | 0.01462 | 0.2013926 |
| Canopy opening | Cano | -0.24649 | 0.0001455 *** |
| Clay content | Clay | 0.20323 | 0.0004671 *** |

\*\*\* significant at 0.1%, \*\* 1%, \* 5% and . 10% probability.

Four models were selected for the richness of the sciophilous species (Table 4), wherein only the magnesium content showed a positive interaction (p < 0.1), while the higher isolation factors of the fragment (p < 0.01), the increase in phosphorus levels (p < 0.1) and the higher slope (p < 0.1) were negative.

**Table 4:** Results of the selected GLM models (ΔAICc < 2) to explain the richness of sciophilous species in the forest fragments of the Guapi-Macacu Basin. The independent variables, the term used in the model, their respective coefficients and their significance by the Chi-square test (χ^2^) are specified below the models.

| Model/Parameters | Term | Coefficient | $\chi^2$ |
| --- | --- | --- | --- |
| Sciophilous = $\beta_0 + \beta_1$ ENN | | $\Delta AICc = 0$ | AICcWi = 0.1614 |
| Intercept | Intercept | 13.81 |  |
| Isolation | ENN | -0.04222 | 0.01327 * |
| Sciophilous = $\beta_0 + \beta_1$ P + $\beta_2$ Mg + $\beta_3$ ENN | | $\Delta AICc = 0.1$ | AICcWi = 0.1522 |
| Intercept | Intercept | 15.42 |  |
| Phosphorous content | P | -60910 | 0.063508 . |
| Magnesium content | Mg | 605.70 | 0.083546 . |
| Isolation | ENN | -0.49 | 0.004779 ** |
| Sciophilous = $\beta_0 + \beta_1$ Mg + $\beta_2$ ENN + $\beta_3$ slope | | $\Delta AICc = 0.2$ | AICcWi = 0.1450 |
| Intercept | Intercept | 14.4 |  |
| Magnesium content | Mg | 13.73 | 0.07294 . |
| Isolation | ENN | -0.04262 | 0.00850 ** |
| Slope | slope | -0.15919 | 0.06733 . |
| Sciophilous = $\beta_0 + \beta_1$ P + $\beta_2$ ENN | | $\Delta AICc = 0.5$ | AICcWi = 0.1241 |
| Intercept | Intercept | 18.06 |  |
| Phosphorous content | P | -0.814630 | 0.165476 |
| Isolation | ENN | -0.04613 | 0.006773 ** |

\*\*\* significant at 0.1%, \*\* 1%, \* 5% and . 10% probability.

The highest weight model (AICcWi = 0.1614) only presented the isolation variable of the fragments (p < 0.05) as explanatory for the species richness of the sciophilous group. In addition to the negative interaction of isolation (p <0.01), the second model (ΔAICc = 0.1; AICcWi = 0.1522) showed negative interaction with the increase in phosphorus content (p < 0.1) and positive interaction with magnesium content (p < 0.1). The third model (AICcWi = 0.1450) added the terrain slope (p < 0.1) as a negative explanation. Finally, the fourth model (AICcWi = 0.1241) only presented isolation (p < 0.01) as a negative factor for sciophyte richness, just as the first model.

## Discussion

### Impacts of intensive agriculture on functional groups

The predictive model results point to a clear influence of the intensive farming environment on the tree community in the forest fragments. This can be observed by the influence that the greater direct limit with agriculture areas has on the abundance of the pioneer species. According to the models, the forest fragments with greater border extension focused on intensive corn and manioc cultivation present greater pioneer species abundance. Furthermore, there seems to be possible indirect impacts caused by the soil eutrophication inside the fragments, possibly due to the aerial drift of fine particles of the soil which are eventually deposited inside these fragments. Thus, the direct limit of a forest fragment with an intensive agriculture area has potential impact on the abundance and richness of the species due to differences in interaction between functional groups and fertility levels, as indicated by the models, where the heliophytes are benefited by calcium levels and disadvantaged by magnesium levels, while the sciophilous species are benefited by magnesium and are disadvantaged by the higher phosphorus levels.

Among the potential impacts from surrounding agricultural areas on the interior of the fragments, changes in soil fertility with increased nutrient levels have been widely discussed in the literature [15,17,18,47]. A recent study in the same study site [18] showed a trend of soil eutrophication in forest fragments located in the same study area, with significant increases in calcium, phosphorus and potassium, while another similar study [17] observed that the higher the fertilizer application intensity in agricultural areas, the higher the nutrient content in nearby forest fragments. The frequent practices of liming and fertilizer use in conventional agriculture fields combined with soil rotation result in drift and transport of fine soil particles by air, which are deposited in soil fragments [15,17, 18]. The result is a significant increase in nutrient levels in the soils of nearby forest fragments such as nitrogen [17,19], phosphorus [15,18], calcium and magnesium [16,18].

The relationship between the abundance and richness of early and late species and the increase in calcium, magnesium and phosphorus nutrient contents found in this study demonstrate that the enrichment of nutrients in the soils of these fragments by the practices implemented in their surroundings is a source of important impact in the tree community and which is still little recognized. This was observed in some studies that identified changes in the floristic composition of herbaceous leaves at the edges of fragments, possibly due to the large input of liming particles for soil acidity correction and the use of nitrogenous fertilizers, causing a dominance of common species in less acidic soils or of nitrophilous species [16,28,29,48,49].

Pioneer tree richness is clearly benefited by increased calcium content, while late species are negatively impacted by increased phosphorus content. These nutrients tend to increase in forest fragments near agriculture areas [16-18] due to the common practice of liming and use of phosphate fertilizers. In addition, phosphorus has low mobility in the soil, which can cause accumulation, and consequently lead to eutrophication of the system over time [18]. Although magnesium content has a positive relationship with the richness of sciophytes, the previous results found by this same study [18] did not indicate significant increases of this nutrient in the fragments, which is the opposite of that found for calcium and phosphorus, as previously mentioned.

Thus, continuous additions of nutrients over long periods may have significant impacts on species composition and functional diversity of the forest community of forest fragments [22] close to conventional agricultural crops, altering the composition and structure of the functional groups of the tree community. Natural ecosystem plants are adapted to conditions with limited availability of nutrients, maintaining a certain balance of adjustment to local fertility characteristics [22]. Thus, soil fertility conditions in tropical forests limit plant growth and establishment, therefore constituting an important determinant of local diversity [19,21,22].

Some recent long-term studies have investigated the primary productivity response of tropical rainforest trees to adding nutrients to the soil [22-25], demonstrating that even in soils relatively rich in exchangeable bases (K^+^, Ca^2+^, Mg^2+^ and Na^+^) and in nitrogen and phosphorus, some species and functional groups respond strongly to the addition of these elements and potassium. However, the impact in regions with lower fertility such as in the areas of this study may be more intense, since these regions tend to have more specialized flora when compared to regions with higher fertility, which have a greater number of fast-growing and more generalist tree species [21,22].

Thus, constant deposition of nutrients in the forest fragment soils of this study may be creating a conducive environment to the proliferation of pioneer tree species. The fragments with an extensive livestock environment showed the highest IV values for the sciophilous species (Table S1), which helps to explain the observed grouping of these fragments in the ordering analysis. Thus, it is possible to observe that the pioneer or secondary heliophilous species may be benefiting in the fragments with an intensive agriculture environment, where the effects of nutrient drift are greater. When analyzing the two functional groups (heliophilous and sciophilous), it is possible to observe that fast growing pioneer species generally present higher rates of primary production in response to nutrient increase [4,50,51]. This is due to the fact that pioneer species seedlings are more efficient in nutrient consumption, being able to stand out in competitive environments with shade tolerant species [51].

However, the environment can affect the composition of the species in two ways: firstly by edge effects, which are more intense when the environment is composed of agriculture, changing the chemical characteristics of the soil and creating direct impacts on the microclimate; and second, by factors that hinder the recruitment of late species by reducing propagule flow [3,4,52–54].

Agriculture areas may also impact late species by influencing the connectivity between the fragments, as they may be a more hostile environment for the movement of pollinators and dispersers [54,55], reducing seed flows [53]. Changes in the number of individuals of some species due to decreased seed dispersal can cause local extinctions over time [53], making it difficult to maintain viable populations. In addition, more intensive use in agricultural areas may create unfavorable microclimatic conditions to the floristic community [56], and may contribute to the impact on the abundance of some shade tolerant species.

### Other impacts on functional groups

The results show that fragment size is one of the factors determining the abundance of species of the two functional groups, as already reported in several studies in tropical forests [2,57-60]. The higher number of sciophilous species was dependent on the fragment size, being the only factor that positively influences the abundance of this group. On the other hand, the abundance of pioneer species is greater in smaller fragments, which is directly related to the fact that these smaller fragments are subject to greater intensity of edge effects [61].

The impact of abrupt edge creation by the fragmentation process for tree communities has been widely discussed, indicating the elevation in mortality rates and changes in the recruitment dynamics of late trees at the edges of fragments [58,62-64], and even in the interior of small fragments [57], benefiting pioneer and generalist species [58,59].

Despite the significant effect on abundance, the species richness for the two functional groups was not affected by the fragment size. For richness, the soil characteristics (fertility levels and clay content), the canopy opening, the slope of the soil and the degree of isolation of the fragments are more important. Plant diversity is highly related to soil fertility and grain size characteristics [19,21,65], and these two factors are strongly correlated as previously discussed. A higher clay content in the soil allows higher amounts of cationic bonds, which provides greater nutrient availability [20,21]. Along these lines, flatter lands have a higher amount of clay coming from higher lands and deposited in these lower areas [20,21], which in turn also implies in higher levels of soil fertility.

In addition to the influence on soil fertility, the slope of the terrain may make it difficult to maintain sciophilous species in two ways; firstly, because the recruitment of these species is difficult as they usually have large seeds which presents greater difficulties to fixate in very sloped areas; secondly, greater slope is associated with higher falling rates and mortality of large trees, since they present greater difficulty to remain fixed in these areas [21]. These situations explain the difficulty in maintaining the seedling recruitment of slow growing species, showing the positive influence of this factor on the abundance of pioneer species, and a negative relation with the abundance and richness of sciophilous species.

The higher degree of isolation of the fragments did not influence the abundance of the species of the two ecological groups, but was one of the explanatory variables for the richness of late species, being more important than the fragment size. As already discussed, this factor leads to reductions in seed flow, decreasing the chances of seed dispersal due to longer distances between fragments [3,4]. Initial successional species have an advantage in seed dispersion because they produce a greater quantity of seeds and generally their seeds have anemochoric dispersion or by generalist animals [34]. On the other hand, late species generally produce larger seeds dispersed by specialist animals [2], making it harder for flows between more isolated fragments.

Finally, unlike what is expected, the larger canopy opening only had a negative influence on the richness of heliophilous species. This result differs from that expected, since greater abundance and diversity of light species are expected for germination and growth in clearings [66]. This unexpected result can be explained by the fact that there was a smaller number of trees with the inclusion diameter (CBH > 15 cm) in the sampled sites where there were clearings due to tree fall, since mortality and the fall of other trees is also generated by the fall of large trees [66], thereby reducing the sampling of pioneers.

Thus, forest fragments immersed in an agricultural matrix and exposed to conventional and intensive agriculture practices may be conditioned to a reduction in tree species diversity through the replacement of late species by pioneer species. Late species are responsible for most of the natural regeneration in mature tropical forests [59], and show a tendency to be restricted to areas with no edge effects reaching them, such as in interiors of large fragments and fragments with little isolation. Thus, the discussion on conservation in agricultural production landscapes should take into account that it is necessary to develop and encourage agricultural practices which are less impacting on biodiversity, in addition to the demand for maintaining large areas of natural remnants which is essential for some species. This is important in the need to reduce the direct impacts caused by edge effects, such as the change in the fertility characteristics of the soils, and the impacts that the matrix quality can imply on the population flows between the fragments [61].

## Conclusion

The study evidences that the intense adoption of chemical inputs for fertilization in agricultural areas causes concrete impacts on the forest community diversity of forest fragments, being more important than the fragment size itself. This study shows that increasing soil fertility of forest fragments adjacent to intensive agriculture areas causes impacts on both abundance and species richness in the ecological groups of the tree community. The increase in phosphorus levels may cause a decrease in the richness of shade tolerant species, while the pioneers benefit from increased calcium levels and the impacts inherent to forest fragmentation such as size reduction and fragment isolation. In this way, a continuous increase in the populations of pioneer species may be causing a regression in the successional stage of these remnants. The ecological intensification of agriculture is a challenge and an emergency for fragmented agricultural landscapes, where the need for biodiversity conservation policies and ecosystem services must be linked to agricultural production. This proves the demand for policies which support developing more conservationist agricultural production strategies, and evidence potential externalities of adopting strategies based on the land-sparing system which are not considered.

## Acknowledgements

We thank all colleagues at the Agricultural Landscapes Ecology Laboratory (*LEPA*) and *EMBRAPA* Agroecology employees for their collaboration in field collection and sample processing. To Daniel Carvalho and all the researchers participating in this research project for the collections and botanical identifications. To Dr. Elaine Fidalgo, from the Geomatics Nucleus (*NGEO*) of *Embrapa* Soils, for obtaining the landscape metrics. We thank the Brazilian Agricultural Research Corporation, *EMBRAPA*-Agrobiology and the Foundation for Research Support of the State of Rio de Janeiro (*FAPERJ*) for funding the research project. Finally, we thank the Commission for the Improvement of Higher Education Personnel (*CAPES*) for providing a scholarship to the first author.

## Supporting Information

**S1 Table. Tree species sampled per strata in forest fragment and their respective basal area values and ecological group classification.**

**S2 Table. Importance Value (IV) matrix of each tree species at each sampling site.**

**S3 Table. Abiotic factors values sampled at each site.** C: carbon; N: nitrogen; P: phosphorus; K: potassium; Ca: calcium; Mg: magnesium; limagri: border limit percentage with agriculture; area: fragment size in hectares; PARA: perimeter-area ratio; ENN: Euclidean distance of the nearest neighbor; Abos: canopy opening; Arg: clay percentage in the soil; decli: slope.

**S4 Table. Tested models for prediction of richness and abundance of sciophilous and heliophilous tree species.**

## References

1. Jackson HB, Fahrig L. Habitat loss and fragmentation. Encycl Biodivers. 2000; 4: 50–58.

2. Laurance WF, Camargo JLC, Luizão RCC, Laurance SG, Pimm SL, Bruna EM, et al. The fate of Amazonian forest fragments: A 32-year investigation. Biol Conserv. 2011; 144(1): 56–67.

3. Magrach A, Laurance WF, Larrinaga AR, Santamaria L. Meta-analysis of the effects of forest fragmentation on interspecific interactions. Conserv Biol. 2014; 28(5): 1342–1348.

4. Moran C, Catterall CP. Responses of seed-dispersing birds to amount of rainforest in the landscape around fragments. Conserv Biol. 2014; 28(2): 551–560.

5. Tilman D, Balzer C, Hill J, Befort BL. Global food demand and the sustainable intensification of agriculture. Proc Natl Acad Sci. 2011; 108(50): 20260–20264.

6. Tscharntke T, Clough Y, Wanger TC, Jackson L, Motzke I, Perfecto I, et al. Global food security, biodiversity conservation and the future of agricultural intensification. Biol Conserv. 2012; 151(1): 53–59.

7. Green RE, Cornell SJ, Scharlemann JPW, Balmford A. Farming and the fate of wild nature. Science. 2005; 307(5709): 550–555.

8. FAO. State of the World’s Forests 2016. Forests and agriculture: land-use challenges and opportunities. Rome; 2016. 107 p. Available from: http://www.fao.org/3/a-i5588e.pdf

9. Foley JA, Ramankutty N, Brauman KA, Cassidy ES, Gerber JS, Johnston M, et al. Solutions for a cultivated planet. Nature. 2011; 478(7369): 337–342.

10. Balmford A, Amano T, Bartlett H, Chadwick D, Collins A, Edwards D, et al. The environmental costs and benefits of high-yield farming. Nat Sustain. 2018; 1(9): 477–85.

11. Perfecto I, Vandermeer J. Biodiversity Conservation in Tropical Agroecosystems. Ann. N.Y. Acad. Sci. 2008; 200: 173–200.

12. Phalan B, Onial M, Balmford A, Green RE. Reconciling Food Production and Biodiversity Conservation: Land Sharing and Land Sparing Compared. Science. 2011; 333: 1289–1291.

13. Tittonell P. Ecological intensification of agriculture-sustainable by nature. Curr Opin Environ Sustain. 2014; 8: 53–61.

14. Fahrig L, Girard J, Duro D, Pasher J, Smith A, Javorek S, et al. Farmlands with smaller crop fields have higher within-field biodiversity. Agric Ecosyst Environ. 2015; 200: 219–234.

15. Duncan DH, Dorrough J, White M, Moxham C. Blowing in the wind? Nutrient enrichment of remnant woodlands in an agricultural landscape. Landsc Ecol. 2008; 23(1): 107–119.

16. Chabrerie O, Jamoneau A, Gallet-Moron E, Decocq G. Maturation of forest edges is constrained by neighbouring agricultural land management. J Veg Sci. 2013; 24(1): 58–69.

17. Didham RK, Barker GM, Bartlam S, Deakin EL, Denmead LH, Fisk LM, et al. Agricultural intensification exacerbates spillover effects on soil biogeochemistry in adjacent forest remnants. PLoS One. 2015; 10(1): 1–32

18. Uzêda MC, Fidalgo ECC, Moreira RV de S, Fontana A, Donagemma GK. Eutrofização de solos e comunidade arbórea em fragmentos de uma paisagem agrícola. Pesqui Agropecu Bras. 2016; 51(9): 1120–1130.

19. John R, Dalling JW, Harms KE, Yavitt JB, Stallard RF, Mirabello M, et al. Soil nutrients influence spatial distributions of tropical tree species. Proc Natl Acad Sci. 2007; 104(3): 864–869.

20. Laurance WF, Fearnside PM, Laurance SG, Delamonica P, Lovejoy TE, Rankin-De Merona JM, et al. Relationship between soils and Amazon forest biomass: A landscape-scale study. For Ecol Manage. 1999; 118: 127–138.

21. Laurance SGW, Laurance WF, Andrade A, Fearnside PM, Harms KE, Vicentini A, et al. Influence of soils and topography on Amazonian tree diversity: A landscape-scale study. J Veg Sci. 2010; 21(1): 96–106.

22. Wright SJ, Yavitt JB, Wurzburger N, Turner BI, Tanner EVJ, Sayer EJ, et al. Potassium, phosphorus, or nitrogen limit root allocation, tree growth, or litter production in a lowland tropical forest. Ecology. 2011; 92(8): 1616–1625.

23. Santiago LS, Wright SJ, Harms KE, Yavitt JB, Korine C, Garcia MN, et al. Tropical tree seedling growth responses to nitrogen, phosphorus and potassium addition. J Ecol. 2012; 100(2): 309–316.

24. Mayor JR, Wright SJ, Turner BL. Species-specific responses of foliar nutrients to long-term nitrogen and phosphorus additions in a lowland tropical forest. J Ecol. 2014; 102(1): 36–44.

25. Alvarez-Clare S, Mack MC. Do foliar, litter, and root nitrogen and phosphorus concentrations reflect nutrient limitation in a lowland tropical wet forest? PLoS One. 2015; 10(4): 1–16.

26. Vitousek PM, Porder S, Houlton BZ, Chadwick OA. Terrestrial phosphorus limitation: Mechanisms, implications, and nitrogen-phosphorus interactions. Ecol Appl. 2010; 20(1): 5–15.

27. Sardans J, Penuelas J. The Role of Plants in the Effects of Global Change on Nutrient Availability and Stoichiometry in the Plant-Soil System. Plant Physiol. 2012; 160(4): 1741–1761.

28. Thimonier A, Dupouey JL, Timbal J. Floristic changes in the herb-layer vegetation of a deciduous forest in the Lorraine Plain under the influence of atmospheric deposition. For Ecol Manage. 1992; 55(1–4): 149–167.

29. Thimonier A, Dupouey Jl, Bost F, Becker M. Simultaneous eutrophication and acidification of a forest ecosystem in North-East France. New Phytol. 1994; 126(3): 533–539.

30. Fidalgo ECC, Pedreira B da CCG, Abreu MB de, Moura IB de, Godoy MDP. Uso e Cobertura da Terra na Bacia Hidrográfica do Rio Guapi-Macacu. Rio de Janeiro: Embrapa Solos. 2008; 32 p. (Embrapa Solos. Documentos 105).

31. Veloso HP, Rangel Filho AL, Lima JCA. Classificação da vegetação brasileira, adaptada a um sistema universal. Rio de Janeiro: IBGE; 1991. 124 p.

32. Laurance WF, Lovejoy TE, Vasconcelos HL, Bruna EM, Didham RK, Stouffer PC, et al. Ecosystem Decay of Amazonian Forest Fragments: a 22-Year Investigation. Conserv Biol. 2002; 16(3): 605–618.

33. Swaine MD, Whitmore TC. On the definition of ecological species groups in tropical rain forests. Vegetatio. 1988; 75(1–2): 81–86.

34. Maciel M de NM, Watzlawick LF, Schoeninger ER, Yamaji FM. Classificação ecológica das espécies arbóreas. Rev Acadêmica ciências agrárias e Ambient. 2003; 1(2): 69–78.

35. Peixoto GL, Martins SV, Silva AF da, Silva E. Composição florística do componente arbóreo de um trecho de Floresta Atlântica na Área de Proteção Ambiental da Serra da Capoeira Grande, Rio de Janeiro, RJ, Brasil. Acta Bot Brasilica. 2004; 18(1): 151–160.

36. Carvalho FA, Nascimento MT, Braga JMA. Composição e riqueza florística do componente arbóreo da Floresta Atlântica submontana na região de Imbaú, Município de Silva Jardim, RJ. Acta Bot Brasilica. 2006; 20(3): 727–740.

37. Sobrinho VG, Schneider PR. Bioeconomic Analysis of Carbon Forest Sequestration and of the Ecological Debt: An Application to the Case of Rio Grande Do Sul. Cienc Florest. 2008; 18(3–4): 493–510.

38. Finotti R, Kurtz BC, Cerqueira R, Garay I. Variations in structure, floristic composition and successional characteristics of forest fragments of the Guapiacu river basin (Guapimirim/Cachoeiras de Macacu, RJ, Brazil). Acta Bot Brasilica. 2012; 26(2): 464–475.

39. Mueller-Dombois D, Ellemberg H. Aims and methods of vegetation ecology. New York: J Wiley; 1974. 547 p.

40. Donagema GK, Campos DVB de, Calderano SB, Teixeira WG, Viana JHM (Org.). Manual de Métodos de Análise de Solo. 2. ed. rev. Rio de Janeiro: EMBRAPA Solos; 2011. 230 p. (EMBRAPA Solos. Documentos 132).

41. Engelbrecht BMJ, Herz HM. Evaluation of different methods to estimate understorey light conditions in tropical forests. J Trop Ecol. 2001; 17(2): 207–224.

42. Frazer, Gordon W, Canham CD, Lertzman KP. Gap Light Analyzer (GLA), Version 2.0: Imaging software to extract canopy structure and gap light transmission indices from true-colour fisheye photographs, users manual and program documentation. Burnaby: Simon Fraser University; Millbrook: Institute of Ecosystem Studies, Copyright © 1999. 36p.

43. Fidalgo ECC, Pedreira, BCCG, Prado RB, Fadul MJ A, Bastos EC, Silva SA, et al. Dinâmica de uso e cobertura da terra das bacias Guapi-Macacu e Caceribu - Relatório e mapa de uso e cobertura da terra das bacias Guapi-Macacu e Caceribu -T0. Contrato Nº 6000.00419115.08.2. [Rio de Janeiro]: EMBRAPA /FAPED. 2011. 18 p.

44. McGarigal K. Fragstats: user guideline. Version 3 [Internet]. Available from: http://www.umass.edu/landeco/research/fragstats/documents/User guidelines/User guidelines content.htm.

45. Dufrene M, Legendre P. Species Assemblages and Indicator Species: The Need for a Flexible Asymmetrical Approach. Ecol Monogr. 1997 Aug; 67(3): 345–366.

46. R. CORE TEAM. R: A language and environment for statistical computing. Viena: R Foundation for Statistical Computing; 2016.

47. Chabrerie O, Jamoneau A, Gallet-Moron E, Decocq G. Maturation of forest edges is constrained by neighbouring agricultural land management. J Veg Sci. 2013; 24(1): 58–69.

48. Boutin C, Jobin B. Intensity of Agricultural Practices and Effects on Adjacent Habitats. Ecol Appl. 1998 May; 8(2): 544–557.

49. Demchik MC, Sharpe WE. Forest floor plant response to lime and fertilizer before and after partial cutting of a northern red oak stand on an extremely acidic soil in Pennsylvania, USA. For Ecol Manage. 2001; 144: 239–244.

50. Baker TR, Swaine MD, Burslem DFRP. Variation in tropical forest growth rates: Combined effects of functional group composition and resource availability. Perspect Plant Ecol Evol Syst. 2003; 6: 21–36.

51. Chapin III FS, M. VP, Van Cleve K. The nature of nutrient limitation in plant communities. Am Nat. 1986; 127(1): 48–58.

52. Gascon C, Lovejoy TE, Bierregaard RO, Malcolm JR, Stouffer PC, Vasconcelos HL, et al. Matrix habitat and species richness in tropcial forest remnants. Biol Conserv. 1999; 91: 223–229.

53. Benitez-Malvido J, Martinez-Ramos M. Influence of edge exposure on tree seedling species recruitment in tropical rain forest fragments. Biotropica. 2003; 35(4): 530–541.

54. Ewers RM, Didham RK. Confounding factors in the detection of species responses to habitat fragmentation. Biol Rev Camb Philos Soc. 2006; 81(1):117–42.

55. Perfecto I, Vandermeer J, Wright A. Nature’s Matrix: Linking agriculture, conservation, and food sovereignty. 1. ed. London: Cromwell Press Group; 2009. 257 p.

56. Mesquita RCG, Delamônica P, Laurance WF. Effect of surrounding vegetation on edge-related tree mortality in Amazonian forest fragments. Biol Conserv. 1999;91(2–3):129–34.

57. Benitez Malvido J. Impact of forest fragmentation on seedling abundance in a tropical rain forest. Conserv Biol. 1998; 12(2): 380–389.

58. Laurance WF, Nascimento HEM, Laurance SG, Andrade A, Ribeiro JELS, Giraldo JP, et al. Rapid decay of tree-community composition in Amazonian forest fragments. Proc Natl Acad Sci U S A. 2006; 103(50): 19010–19014.

59. Santos BA, Peres CA, Oliveira MA, Grillo A, Alves-Costa CP, Tabarelli M. Drastic erosion in functional attributes of tree assemblages in Atlantic forest fragments of northeastern Brazil. Biol Conserv. 2008; 141(1): 249–60.

60. Munguía-Rosas MA, Montiel S. Patch size and isolation predict plant species density in a naturally fragmented forest. PLoS One. 2014; 9(10).

61. Laurance WF, Vasconcelos HL. Consequências ecológicas da fragmentação florestal na amazônia. Oecologia Bras. 2009; 13(03): 434–451.

62. Laurance WF, Laurance SG, Ferreira LV, Merona, Judy MR, Gascon C, Lovejoy TE. Biomass Collapse in Amazonian Forest Fragments. Science. 1997; 278(5340): 1117–1118.

63. Mesquita RCG, Delamônica P, Laurance WF. Effect of surrounding vegetation on edge-related tree mortality in Amazonian forest fragments. Biol Conserv. 1999; 91(2–3): 129–134.

64. Laurance WF, Ferreira L V., Merona JMR, Laurance SG. Rain Forest Fragmentation and the Dynamics of Amazonian Tree Communities. Ecology. 1998 Sep; 79(6): 2032.

65. Oliveira-Filho AT, Curi N, Vilela EA, Carvalho D. Variation in Tree Community Composition and Structure With Changes in Soil Properties Within a Fragment of Semideciduous Forest in South-Eastern Brazil. Edinburgh J Bot. 2001; 58(1): 139–158.

66. Martins SV. Sucessão Ecológica: Fundamentos e aplicações na restauração de ecossistemas florestais. In: Martins SV (Ed). Ecologia de Florestas Tropicais do Brasil. 2. ed. Viçosa: Editora UFV; 2012. p. 372.

